# A machine-learning-guided mutagenesis platform for accelerated discovery of novel functional proteins

**DOI:** 10.1101/282996

**Authors:** Yutaka Saito, Misaki Oikawa, Hikaru Nakazawa, Teppei Niide, Tomoshi Kameda, Koji Tsuda, Mitsuo Umetsu

## Abstract

Molecular evolution based on mutagenesis is widely used in protein engineering. However, optimal proteins are often difficult to obtain due to a large sequence space that requires high costs for screening experiments. Here, we propose a novel approach that combines molecular evolution with machine learning. In this approach, we conduct two rounds of mutagenesis where an initial library of protein variants is used to train a machine-learning model to guide mutagenesis for the second-round library. This enables to prepare a small library suited for screening experiments with high enrichment of functional proteins. We demonstrated a proof-of-concept of our approach by altering the reference green fluorescent protein (GFP) so that its fluorescence is changed to yellow while improving its fluorescence intensity. Using 155 and 78 variants for the initial and the second-round libraries, respectively, we successfully obtained a number of proteins showing yellow fluorescence, 12 of which had better fluorescence performance than the reference yellow fluorescent protein (YFP). These results show the potential of our approach as a powerful platform for accelerated discovery of functional proteins.

## Introduction

Molecular evolution based on mutagenesis is widely used in protein engineering, where critical amino acid residues of a target protein are identified based on available structural information and mutated for function alteration and maturation. Given a number of critical positions *k*, there are 20^*k*^ possible sequences. In iterative saturation mutagenesis (ISM), one of the principal molecular evolution methods, mutagenesis proceeds in a step-wise manner (*1-3*): a residue is mutated in all possible ways and the optimal residue is selected through experimental evaluation. The residue is fixed and the next residue is determined in the same manner. Since the effects of mutations on function are often synergistic or antagonistic, ISM does not always lead to the optimal sequence. On the other hand, the library approach mutates all critical residues simultaneously via evolution operations and allows us to discover optimal sequences under synergistic or antagonistic coupling (*4, 5*). Recent advances in genetic engineering allowed us to prepare an extremely large library, beyond the limit of organic synthesis (*6-8*). Such a large library, however, leads to high costs in screening experiments. The success of protein engineering crucially depends on preparing a small library with high enrichment of functional proteins.

In this study, we propose a novel approach that combines molecular evolution with machine learning to accelerate the discovery of functional proteins. In this approach, a Gaussian process is trained with an initial small library to propose the second-round mutagenesis library. The proteins in the second-round library are chosen according to the probability-of-improvement acquisition function commonly used in Bayesian optimization. We show the potential of this approach for efficiently altering protein function, by driving fluorescence color change in the green fluorescence protein (GFP) (*9*). A small library of GFP variants was generated by means of point saturation and site-directed random mutagenesis. Sequence and functional data acquired from the variants in the library were used for training a machine-learning model to create the second-round library. Mix primer techniques in overlap extension polymerase chain reaction (PCR) generated the second-round library which consisted of ∼80 variants proposed by machine learning. The library turned out to contain yellow fluorescent proteins (YFPs) in high number. These results show the potential of our approach as a powerful platform for accelerated discovery of functional proteins.

## Results

### Overview of the platform

The overview of our platform is shown in Fig. 1. The workflow starts from a target protein whose function is to be improved or altered. First, an initial library of protein variants is generated by means of conventional mutagenesis approaches, such as point saturation mutagenesis and random mutagenesis. Some of the generated variants (typically between tens and hundreds) are prepared in small-scale bacterial cultivation with a deep-well plate. Data about gene sequences, concentration, and performance (e.g., fluorescence) of the variants are quickly obtained from the bacterial lysates, using a DNA sequencer and a plate reader in combination with the suitable assay. This data is used to train a Bayesian machine-learning model such as Gaussian processes. The trained model is used to rank all possible variants according to the probability of having desirable functions. The second-round library of top-ranked variants is prepared from the gene fragment of the original protein by using a mixture of primers with designed mutagenesis codons. The data from the second-round library is then in turn acquired. Thanks to the enhancement provided by the machine-learning approach, the best proteins in the second-round library are typically much better than those in the initial library.

**Fig. 1.**
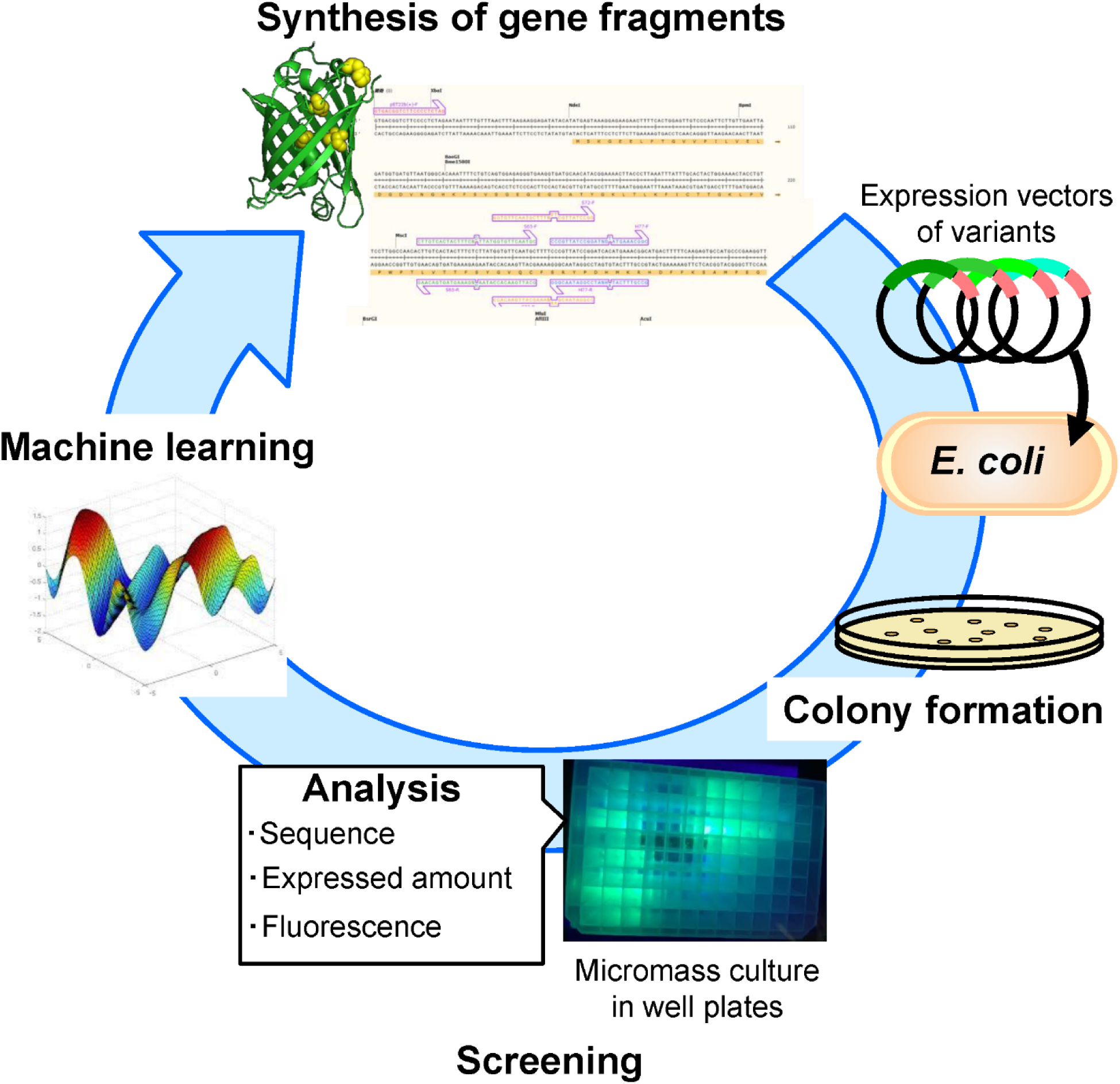
Overview of the machine-learning-guided mutagenesis platform for evolving GFP to YFP.

### Preparation of the initial data of GFP variants for machine learning

In this study, we present an application of our platform where the cycle3 GFP (*9*) is altered so that its fluorescence is changed from green to yellow, and its fluorescence intensity is improved. We surveyed existing GFPs and YFPs (*10-14*), and a new YFP, referred to as “reference YFP”, was prepared by introducing four mutations (S65G, S72A, H77Y, and T203F) in the cycle3 GFP (Table S1). An initial library of GFP variants was obtained by applying two different methods: point saturation mutagenesis and site-directed random mutagenesis, to these residues.

Fig. 2A shows yellow fluorescence ratio and maximum fluorescence intensity of the variants obtained from point saturation mutagenesis. Here, saturation mutagenesis by means of 22-c trick method (*15*) was independently applied to each residue, to make saturated groups of GFP variants where only one of the four residues is mutated (i.e., 19 × 4 = 76 variants). The gene fragments of GFP variants with saturated mutagenesis at one residue were amplified by PCR with a pair of designed 22-c trick primer (Table S2), and the mixed gene fragments were ligated in expression vectors in one pot (*16*). *Escherichia* (*E.*) *coli* bacteria were transformed with the mixture of the ligated vectors and they were spread on agar culture plates to form colonies, each of which should contain a vector bearing a gene fragment. The picked colonies were separately grown in a well of 96-deep-well plates. From each cell lysate, GFP variants were identified and their fluorescence spectra were measured. Upon measuring, 40 out of 76 variants were fluorescence-active in terms of maximum fluorescence intensity over 200. Among them, S72- or H77-mutated variants did not show large positive change in fluorescence intensity nor yellow fluorescence ratio compared to the reference GFP. On the other hand, the mutations at S65 and at T203 critically influenced fluorescence intensity and yellow fluorescence ratio, respectively (Fig. 2A). From these S65- or T203-mutated variants, 11 variants were fluorescence-active, and selected as training data for machine learning.

**Fig. 2.**
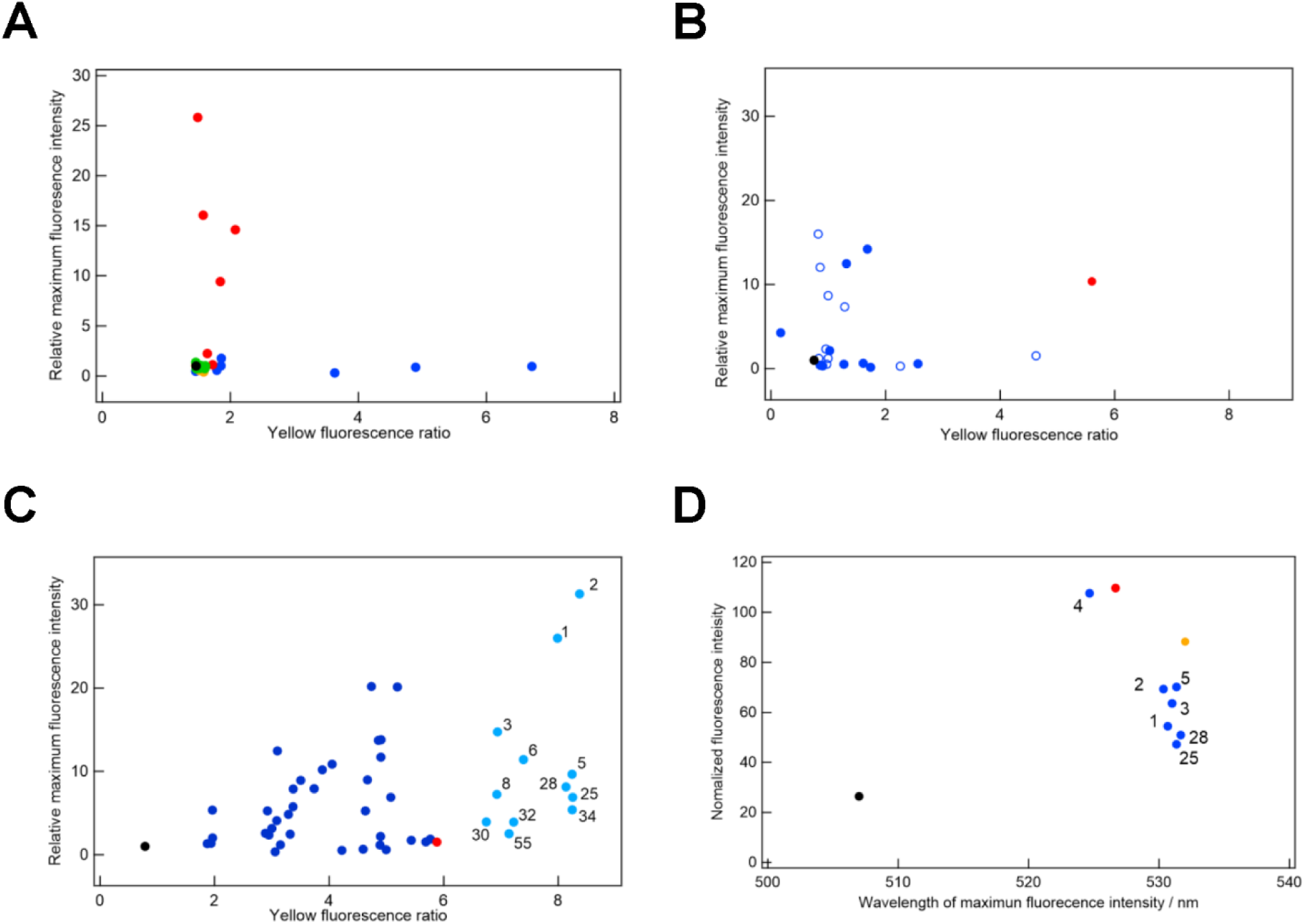
Fluorescence properties of GFP variants. (**A**) Saturation mutagenesis was conducted at S65 (red circles), S72 (orange circles), H77 (green circles), and T203 (blue circles). The black circle represents the reference GFP. (**B**) Saturation mutagenesis (open blue circles) and random mutagenesis (closed blue circles). Black and red closed circles represent the reference GFP and YFP, respectively. (**C**) GFP variants proposed by machine learning (blue and light blue circles). Light blue circles represent 12 variants with yellow fluorescence ratio higher than the reference YFP. The number represents the ranking by machine learning. Black and red circles represent the reference GFP and YFP, respectively. (**D**) Fluorescence properties of the purified GFP variants. Blue circles represent variants proposed by machine learning with the numbers showing their ranks. Black, red, and orange circles represent the reference GFP, the reference YFP, and Venus, respectively.

In site-directed random mutagenesis, the four residues were simultaneously randomized and 142 GFP variants were obtained. The gene fragments of the variants were produced by PCR with a mixture of the 22-c trick primers (Table S2). The pairs of the 22-c trick primers for S65, H77, and T203 were simultaneously used in PCR, and the amplified fragments were amplified again with the pair of the 22-c trick primers for S72. After the ligation of the amplified gene fragments into the opened expression vectors, *E. coli* transformed with the vectors were grown on agar culture plates and then in a 96-deep-well plate. 186 colonies were picked up, and their GFP variants were sequenced together with a measurement of the fluorescence activity. Consequently, 142 GFP variants were identified, among which 10 were fluorescence-active.

In total, we generated 218 variants (76 by point saturation mutagenesis plus 142 by site-directed random mutagenesis) as well as the reference GFP and YFP. However, no variant had better fluorescence properties compared to the reference YFP (Fig. 2B). This result highlights the difficulty of obtaining high-performance variants from purely random exploration.

### Machine learning with the initial library

To create the second-round library, we constructed a machine-learning model that predicts the fluorescence performance of GFP variants from their amino acid sequences. The performance score is defined so that it takes a large value only if both the fluorescence intensity and the yellow fluorescence ratio are high (Materials and Methods).

A Gaussian process model was trained using 155 proteins from the initial library (Table 1). They included 142 variants prepared by site-directed random mutagenesis (10 fluorescent and 132 non-fluorescent variants) together with the reference GFP and YFP. In addition, 11 fluorescent S65- or T203-mutated variants prepared by point saturation mutagenesis were also included in training data to increase the number of fluorescent variants.

**Table 1.**
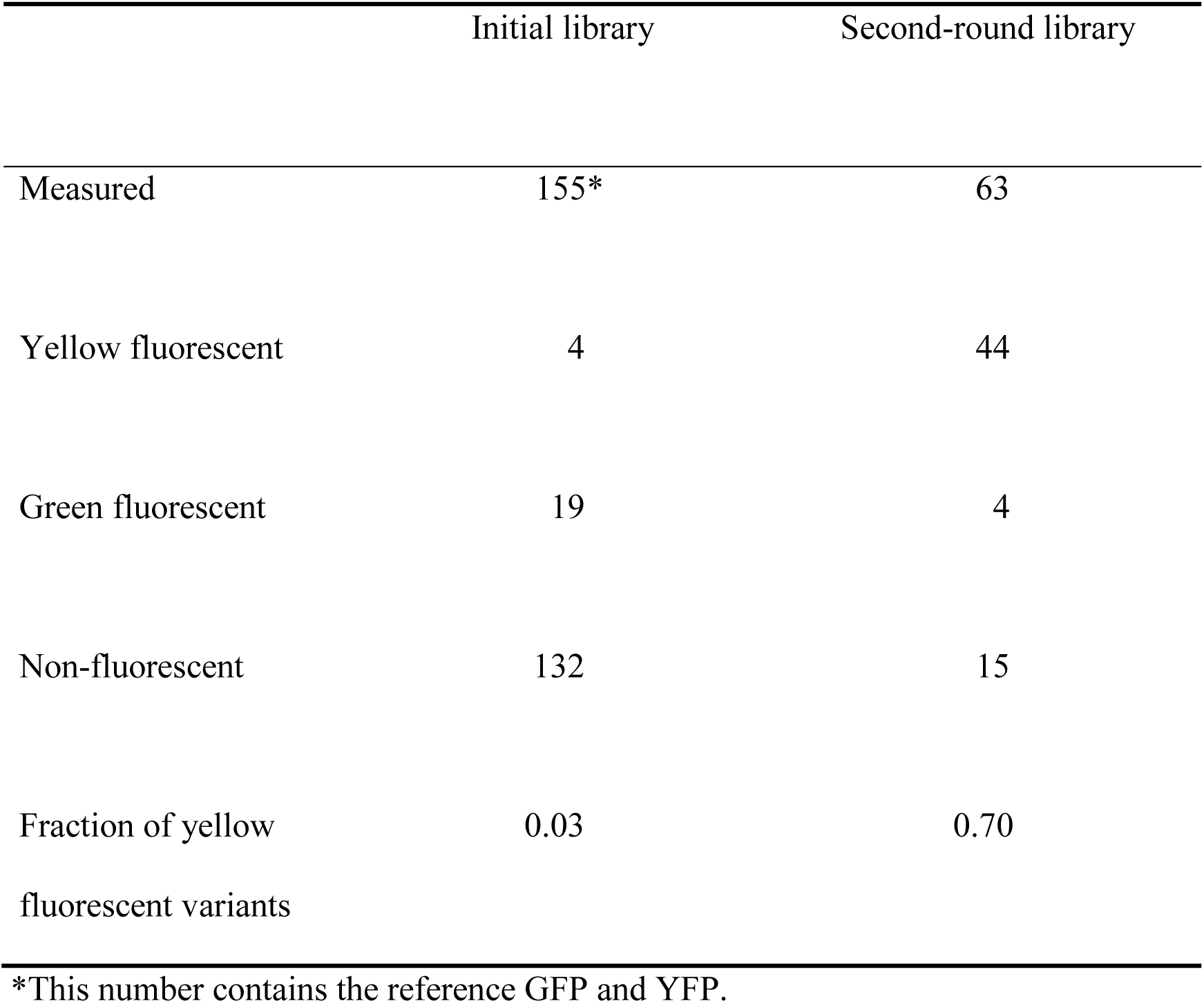
Summary of the fluorescence properties of GFP variants. GFP variants in the initial library used for machine learning, and those in the second-round library proposed by machine learning are shown.

We considered a variety of amino acid descriptors based on physicochemical properties or structural topology (Materials and Methods) and found that the T-scale descriptor (*17*) achieves the best accuracy for our problem by benchmark experiments. Using the trained model, we ranked all unknown variants in the sequence space using the probability-of-improvement score (Data file S1). Interestingly, we found that the second-ranked variant had the same amino acids at the four mutated residues (S65G, S72A, and T203Y with H77 unchanged) as Venus (*18*): an enhanced YFP previously discovered by a conventional mutagenesis approach. The remaining high-ranked variants were previously unknown to the best of our knowledge.

### Construction of a machine-learning-guided mutagenesis library

To prepare the second-round library of GFP variants, we selected the top 78 variants proposed by machine learning (Table 2). The reason of this choice was two-fold: first, the library size of 78 is small and suited for screening experiments; second, the library of these 78 variants can be prepared so that it contains non-candidate variants (i.e., those not proposed by machine learning) at a minimal fraction, by using the following method. The 78 candidate variants were grouped into five classes based on the mutagenesis codons for S65, S72, and H77 (Table 3), which should be contained in mutagenesis primers (Table S3). The pair of the mutagenesis primers in each class was used to amplify the gene fragments from four expression vectors where the GFP variants had T203 mutated to F, H, W, and Y. The prepared library contained only three non-candidate variants besides the 78 candidate variants (Table 3).

**Table 2.**
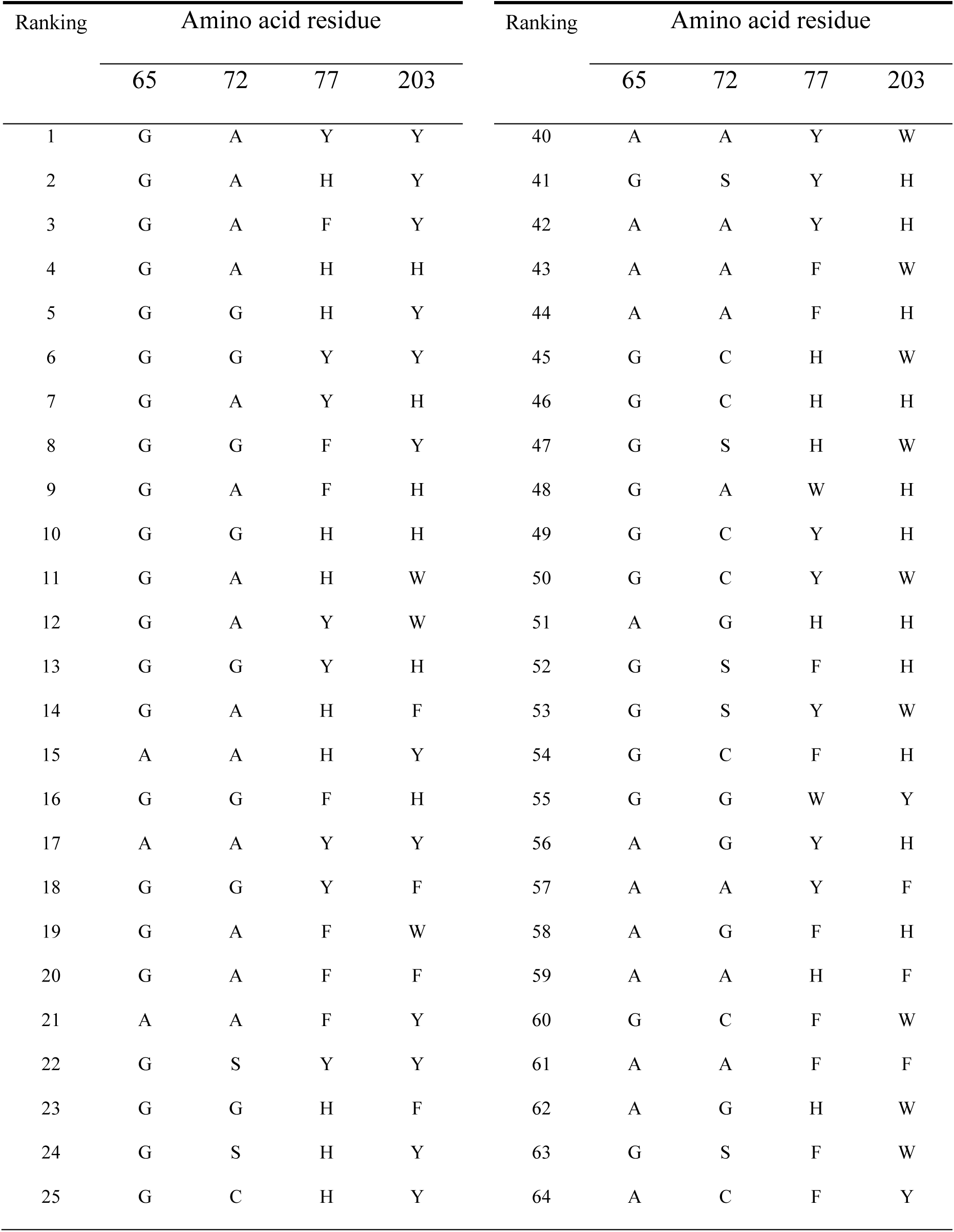
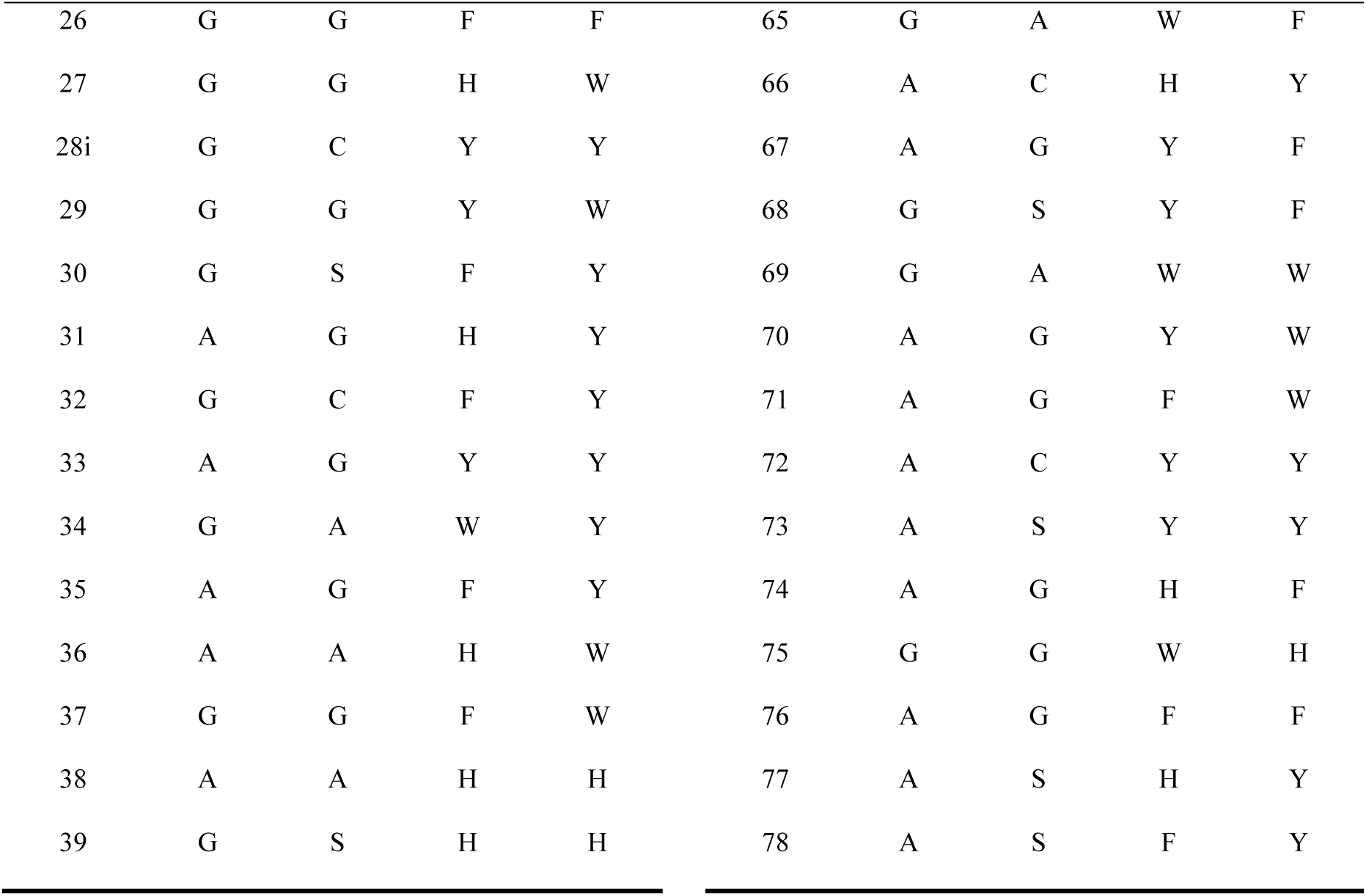
The top 78 candidate variants ranked by machine learning. The complete ranking of all possible variants is given in Data file S1.

**Table 3.**
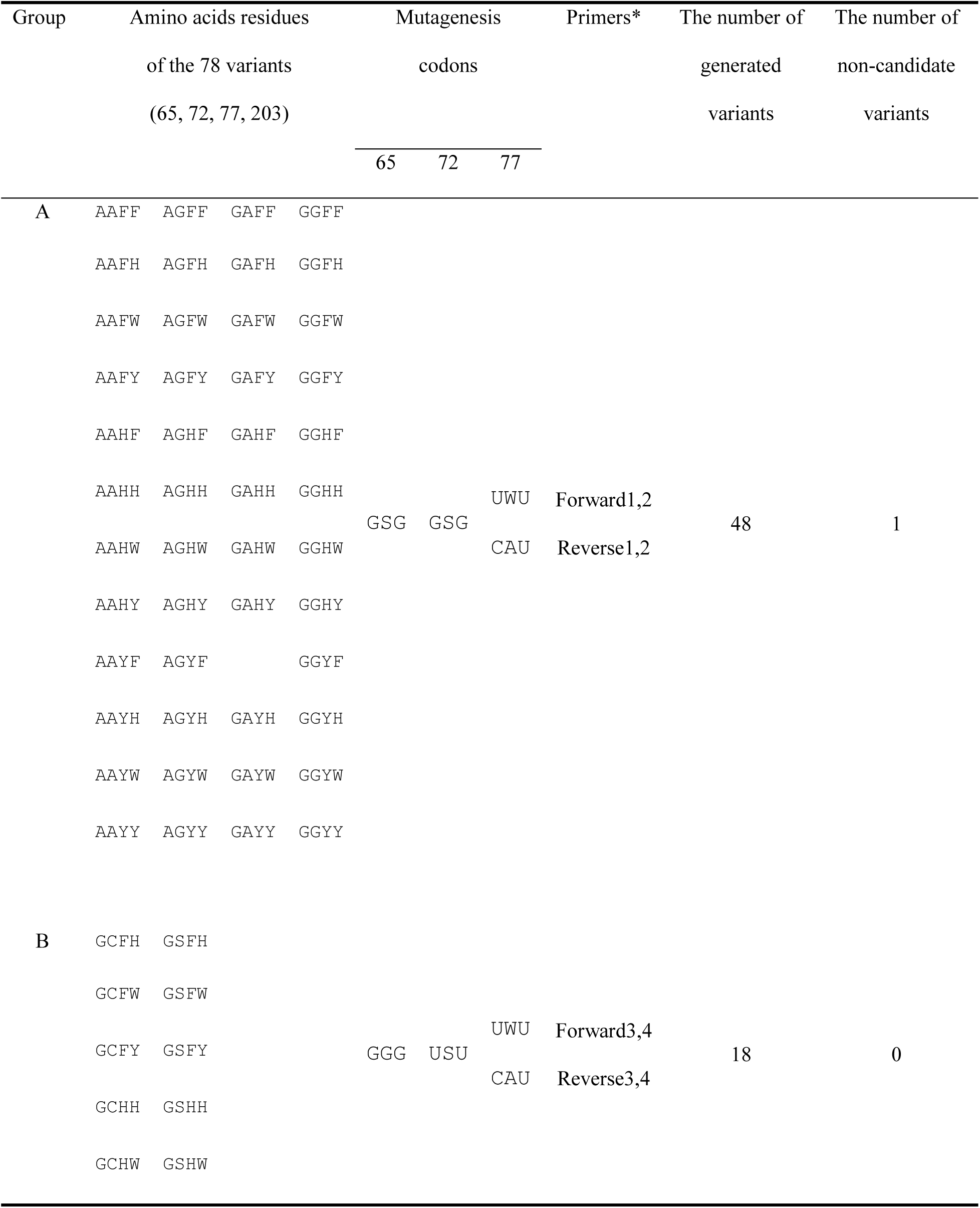
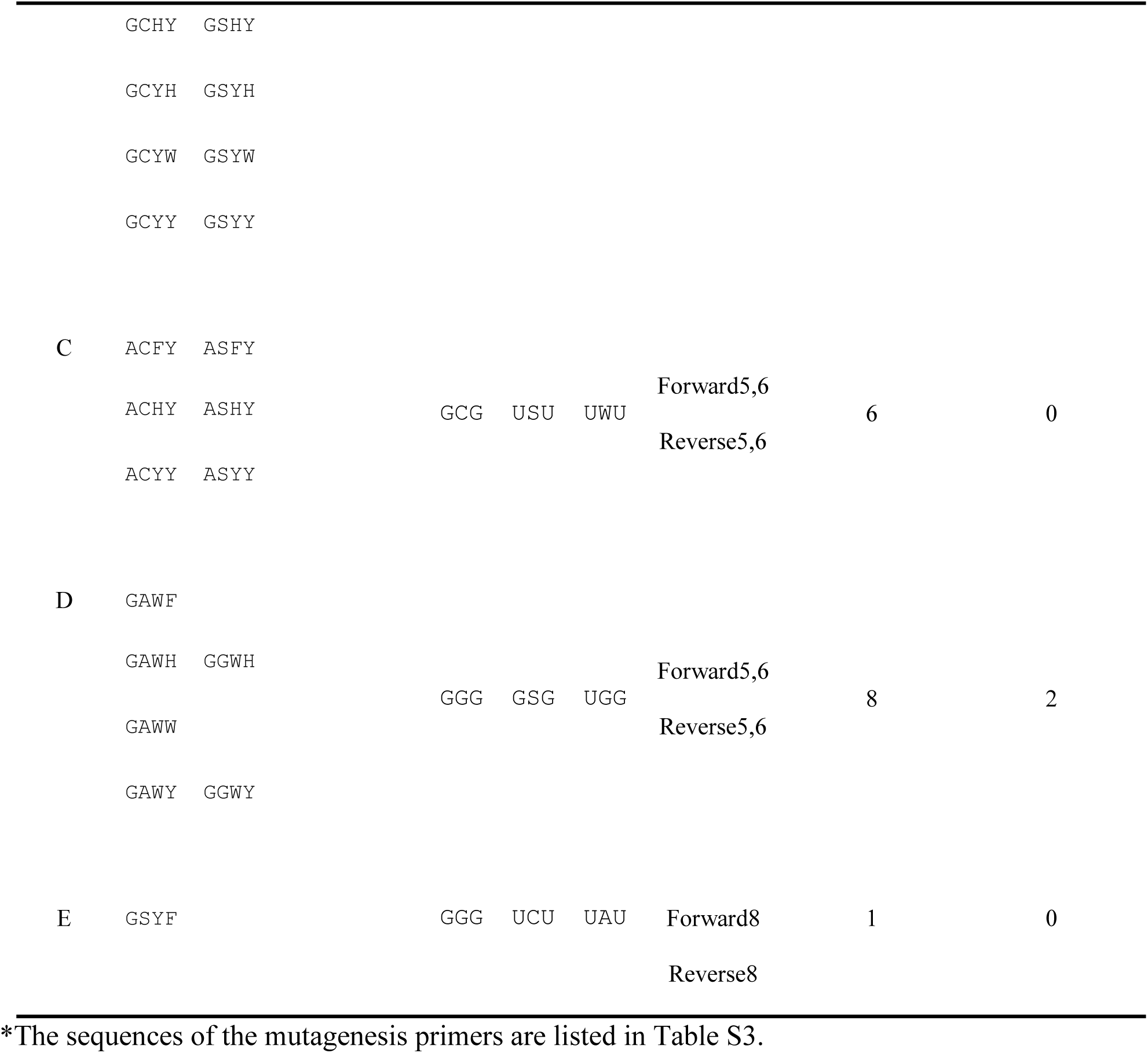
Mutagenesis codons for the second-round library of the top 78 candidate variants proposed by machine learning.

### Screening the GFP variants in the second-round library

In the same way as the initial library, the amplified fragments with the mutagenesis primers listed in Table S3 were ligated into the opened expression vectors, and the *E. coli* transformed with the vectors was grown to form colonies. Eventually, 63 out of the 78 candidate variants were identified from 352 colonies (Table S4), and their yellow fluorescence ratio and maximum fluorescence intensity were measured (Fig. 2C). Surprisingly, the fraction of variants showing yellow fluorescence was much higher than the first library (Table 1), and 12 variants had a yellow fluorescence ratio higher than the reference YFP (light blue circles in Fig. 2C). Moreover, most of the top five candidate variants, except for the 4th-place variant, belonged to the group of these high-performance variants. This result illustrates the effectiveness of machine learning to guide mutagenesis for improved fluorescence performance.

The top five candidate variants and the two variants (the 25th- and the 28th-place variants in Fig. 2C) with the comparable yellow fluorescence ratio were purified by affinity and size exclusion chromatography to quantify their fluorescence intensity and their wavelength at the maximum fluorescence intensity (Fig. 2D). Except for the 4th-place variant, the fluorescence of all variants was red-shifted compared to the reference YFP, achieving the wavelengths similar to Venus. Machine learning demonstrably guided the evolution of GFPs towards high-performance YFPs. The 4th-place variant had a blue-shifted fluorescence compared to the other variants; its wavelength at the maximum fluorescence intensity was similar to that of the reference YFP whereas its fluorescence intensity was the highest among the other variants.

We note that the yellow fluorescence ratio measured from the purified variants (Fig. 2D) were consistent with those measured in the screening assay (Fig. 2C), while some difference was observed regarding the fluorescence intensity. It is well known that fluorescent proteins are sensitive to solution conditions such as pH and ions (*19, 20*). The fluorescence in the screening assay was measured in the lysate solution containing some detergents whose components are not disclosed. The observed variance of the fluorescence intensity thus might be attributed to the difference of pH and components.

## Discussion

### Roles of mutations in yellow fluorescence

Our machine-learning-guided mutagenesis platform found a number of protein variants whose fluorescence performance is better than or comparable to the reference YFP. Analysis of these variants enables us to gain insights into the principles underlying yellow fluorescence. Among the 12 variants with better fluorescence performance than the reference YFP, we noticed several characteristic aspects of advantageous mutations at each residue (Fig. 3).

**Fig. 3.**
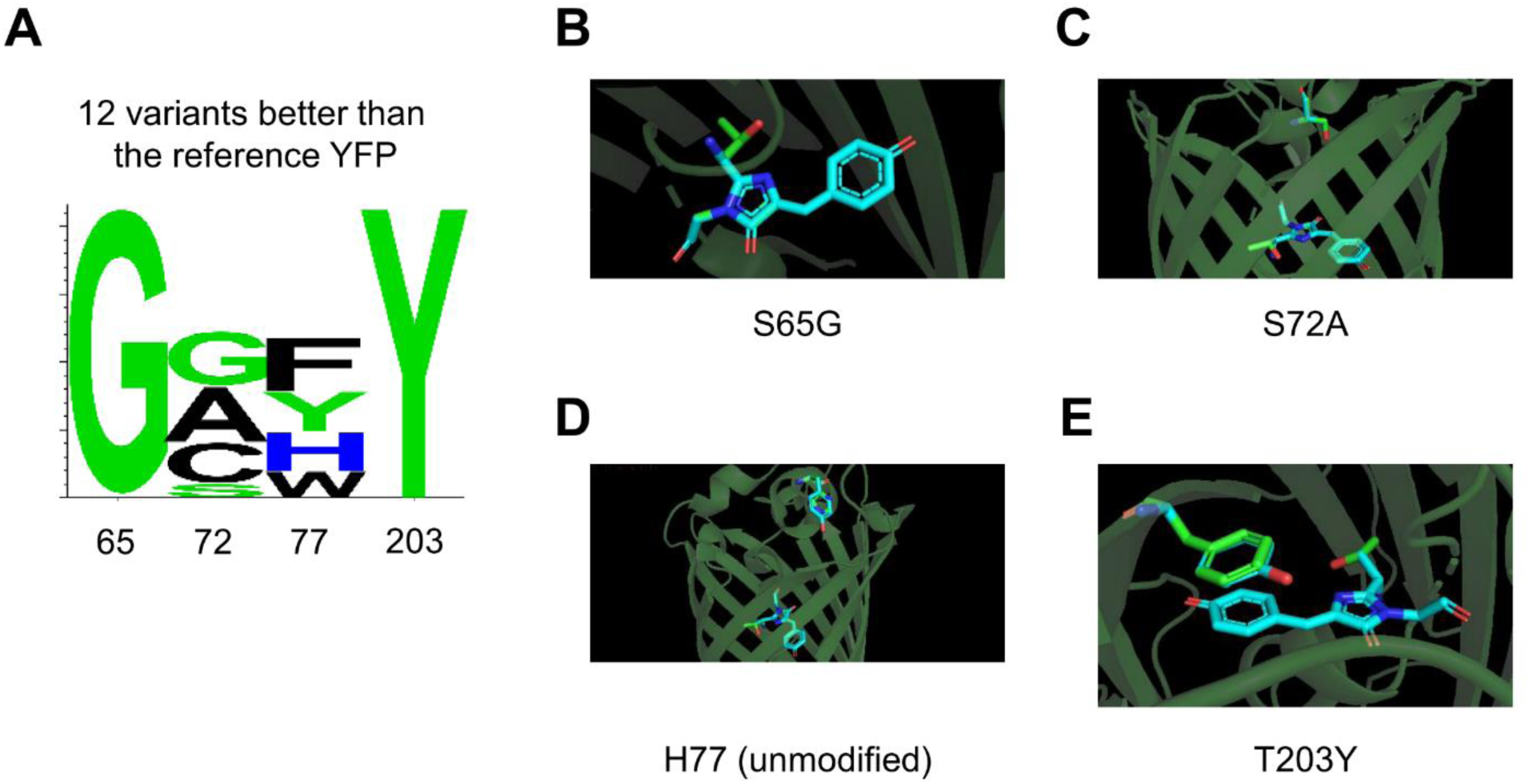
Analysis of 12 protein variants showing higher fluorescence performance than the reference YFP. (**A**) Mutation profile at each residue shown by a sequence logo (*37*). (**B-E**) Overlay of the trace of the reference GFP (green) and a representative variant (cyan) with the mutations (**B**) S65G, (**C**) S72A, (**D**) H77 (unmodified), and (**E**) T203Y. The chromophore (corresponding to the residues 65-67) is also highlighted. As the reference structure, the cycle3 GFP is used (PDB ID code: 2B3P). The structure is drawn by using PyMOL (*38*).

For example, S65 was mutated to G in all of the 12 variants, suggesting a strong advantage of the mutation at this residue (Fig. 3AB). This is consistent with previous studies (*21, 22*) that have reported that the mutation to small aliphatic amino acids such as A, C, L, T, and G induces a red-shifted excitation spectrum and improves molar extinction, possibly due to the ionization of E222 by changing the hydrogen-bonding network around the chromophore, and the complete ionization of Y66. Interestingly, among these small aliphatic amino acids, the mutation to G has resulted in the most red-shifted variant, which was consistent with our results (Fig. 3A). In a similar situation, at T203, the mutation to Y was observed in all of the 12 variants (Fig. 3AE). It has been known that aromatic mutations at T203 produce the π-π interaction between the aromatic ring of the residue 203 and the chromophore phenol ring, which reduces the excited state energy, thereby increasing the excitation and emission wavelengths (*13, 23*). Experimentally, all aromatic amino acids tested have caused a shift in excitation and emission spectra in the order of Y > F > H > W. Our prediction of the mutation at T203 is thus in an agreement with the fact that T203Y creates the biggest red shift.

In contrast to S65G and T203Y, S72 showed a more relaxed type of mutations, accepting the mutation to either of A, G, or C (Fig. 3AC). The mutation of S72A is known to enhance the folding efficiency, leading to high brightness at relatively high temperature (*24, 25*). Similarly, the brightness of the S72G variant has been reported to be higher than that of the wild-type GFP (*26*). However, there is no report about S72C as far as we know. These amino acids (A, G, and C) have a common property that the sizes (i.e., van der Waals volume) of the side chain are small compared to other amino acids, whereas their hydrophobicity is diverse (A and G are hydrophobic, while C is hydrophilic). Thus, for the evolution of S72, the size of the side chain is more important than other features and may affect the stability of GFP. In some of the 12 variants, H77 remained unmodified, while others accepted the mutation to Y, F, or W (Fig. 3AD). A common property of H, Y, F, and W is that they are aromatic amino acids. However, W appeared only in two of the 12 variants, probably due to its large volume relative to H, Y, and F. Therefore, at this site, aromatic amino acids with a moderate volume may have a beneficial effect for yellow fluorescence.

In summary, these analyses show the potential of our platform not only for accelerated discovery of functional proteins, but also as a means of suggesting the roles of mutations underlying molecular evolution.

### Combination of molecular evolution techniques and machine learning

From the initial library generated by point saturation and site-directed random mutagenesis, we could not find any protein variants showing higher fluorescence performance than the reference YFP (Fig. 2B); especially, in site-directed mutagenesis, 132 out of 155 variants showed no detectable fluorescent activities (Table 1). These results highlight the limitation of conventional mutagenesis approaches in finding novel proteins with desirable functional properties. In contrast, our machine-learning method succeeded in selecting high-performance fluorescent proteins from all possible variants. Indeed, the second-round library generated by machine-learning-directed mutagenesis was substantially enriched with high-quality variants (Table 1). Moreover, 12 out of 63 synthesized variants achieved better fluorescence performance than the reference YFP (Fig. 2C). These included a variant strongly similar to Venus (*18*) as well as previously-unknown yellow fluorescent proteins, demonstrating the potential of our machine-learning-guided mutagenesis platform.

While our platform was applied to GFP in this study, it can be used to explore mutagenesis design for various functional alternation of proteins, such as catalytic and thermotolerant functions. For this purpose, fluorescence assays applied in this study can be changed to other measurements depending on the function of proteins to be altered. In addition, machine-learning methods may use different amino acid descriptors and/or other types of feature values, e.g., those calculated by molecular dynamics simulation. These points need to be studied as a future direction.

## Materials and Methods

### Preparation of gene fragments of GFP variants

For point saturation mutagenesis at each residue of S65, S72, H77, and T203, the 22-c trick method was employed (*15*). The gene fragments coding the GFP variants where one of the residues was mutated were generated from the plasmid containing the cycle3 GFP (*9*) by means of overlap extension PCR (*27*), using the external and 22-c trick primers shown in Table S2.

For site-directed random mutagenesis, the three residues of S65, H77, and T203 were simultaneously mutated by means of overlap extension PCR with the external primers and the 22-c trick primers for S65, H77, and T203. The amplified fragments were purified by means of gel extraction, and they were amplified again by means of overlap extension PCR with the 22-c trick primers for T203.

To generate the mutagenesis library guided by machine learning, the gene fragments of GFP variants were amplified from the T203F, T203H, T203W, and T203Y GFP variants, in the overlap extension PCR with the pairs of mutagenesis primers in each class shown in Table S3.

### Preparation of GFP variants

The amplified gene fragments of GFP variants were digested with *Not*I and *Nde*I, then were ligated into the linear (*Not*I- and *Nde*I-digested) pET22b vectors. *E. coli* bacteria were transformed with the resultant vectors and spread on an agar media plate containing 100 μg/mL ampicillin to form colonies (*16*). The colonies grown on the agar media plates were randomly picked up and incubated overnight in 1 mL of LB broth containing 100 μg/mL ampicillin with a deep-well plate (Axygen, CA, USA). 1 μL of each cell culture was used for analyzing gene sequences and another 100 μL was inoculated into 900 μL of the 2×YT broth supplemented with 100 μg/mL ampicillin in a deep-well plate. After incubating for 3 h at 37 °C, isopropyl-1-thio-L-D-galactopyranoside was added to each well to a final concentration of 1 mM to induce the GFP variant expression, and cells were incubated for a further 6 h. The harvested cells were collected and then 200 μL of BugBuster solution (MerK, NJ, USA) was added. The suspensions were incubated for 30 min at 4 °C, and the lysates were centrifuged to remove insoluble matters to measure fluorescence spectra of GFP variants in the supernatants.

### Gene sequence analysis and fluorescence measurement for GFP variants

1 μL of cell culture was mixed with 19 μL of PCR solution containing Ex taq DNA polymerase, and the gene sequences of the amplified fragments by PCR with T7 pro/term primers were analyzed with a 3130 *xl* Genetic Analyzer (Applied Biosystems Inc., CA, USA). Fluorescence spectra from the lysates from transformed *E. coli* were measured with a Synergy H4 hybrid multi-mode microplate reader (BioTek Japan, Tokyo, Japan) at the excitation wavelength of 475 nm. The fluorescence activity of GFP variants were estimated from normalized fluorescence intensity where the intensity is divided by GFP concentration estimated by tag-sandwich ELISA, and yellow fluorescence ratio where total of average fluorescence intensities from 503 nm to 509 nm and from 523 nm to 527 nm is divided by the first intensity (503-509 nm).

### Sandwich ELISA

50 μL of 2 μg/mL anti-GFP antibodies in phosphate buffered saline (PBS) were incubated in the wells of a 96-well polystyrene ELISA microplate (655061; Greiner, Austria) for 1 h, and then 150 μL of 5 w/v% bovine serum albumin in PBS was added to the wells. After washing each well with 0.005% Tween 20 in PBS, 27-fold diluted lysates were incubated for 30 min. The wells were washed again with 0.005% Tween 20 in PBS, and then incubated for 40 min at 25 °C with a 2500-fold dilution of a commercial solution of horseradish peroxidase (HRP)-conjugated anti-His-tag antibody (final concentration, 1.3 nM; sc-8036; Santa Cruz Biotechnology, TX, USA). After washing with 0.005% Tween 20 in PBS, 50 μL of 3,3’,5,5’-tetramethylbenzidine solution (1-step Ultra TMB-ELISA, Thermo scientific, MA, USA) was added and incubated for 2-10 min at 37 °C, and then absorbance at 450 nm was measured after the addition of 50 μL of 2 M H_2_SO_4_.

### Machine learning model

We used a machine learning method based on COMBO (*28*), a fast implementation of Bayesian optimization (*29*) that we have previously developed for material science. Briefly, COMBO uses a Gaussian process based on a liner regression model with random feature map:

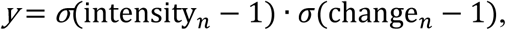

where *y* is the fluorescence performance score of the protein (defined in the next section), ***x*** is a feature vector of the protein, ***φ(x)*** is a random feature map from ***x*** to a ***d***-dimensional numerical vector (***d*** = 5000 in this study), and *ε*is an error term. Given a set of training data {(y, x)}, COMBO fits a ***d***-dimensional weight vector ***w*** so that the fluorescence performance score *y* can be predicted from the feature vector ***x***. For each unknown protein not included in the training data, COMBO can evaluate the probability-of-improvement score (*29*) that represents the probability that the fluorescence performance of the protein is higher than any measured proteins in the training data. These values were used to rank unknown proteins in the space of all possible amino acid sequences (Data file S1).

### Fluorescence performance score

The fluorescence performance of each measured protein was evaluated based on its fluorescence intensity and yellow fluorescence ratio. Since COMBO does not support the simultaneous optimization of multiple properties, we combined these two measures into a single fluorescence performance score:

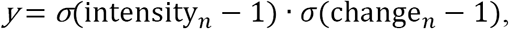

where *σ*(·) is a sigmoidal function, intensity_*n*_ is the fluorescence intensity of the protein divided by that of the reference GFP, change_*n*_ is the yellow fluorescence ratio of the protein divided by that of the reference GFP. This score takes a high value only if both the yellow fluorescence ratio and the fluorescence intensity are high, enabling COMBO to optimize both of the two properties.

### Feature vector

As a feature vector ***x*** for the protein, we used a pre-computed feature vectors for amino acids. Specifically, we defined a feature vector of a GFP variant by concatenating the feature vectors of amino acids at the four mutated sites. For a feature vector of each amino acid, we tested a variety of amino acid descriptors including Z-scale (*30*), T-scale (*17*), ST-scale (*31*), FASGAI (*32*), MS-WHIM (*33*), ProtFP (*34*), VHSE (*35*), and BLOSUM-based features proposed in (*36*). We compared the effectiveness of these descriptors by a benchmark experiment where COMBO was set to find the reference YFP from the initial library using a Bayesian optimization procedure (Fig. S1). In this experiment, Z-scale, T-scale, and ST-scale achieved better results than other descriptors in terms of the number of training rounds for finding the reference YFP. These three descriptors were further compared in another benchmark experiment where COMBO was trained on the initial library with the reference YFP excluded, and all possible variants in the sequence space of 20^4^ were ranked by probability-of-improvement scores. In this experiment, T-scale achieved the best result in terms of the rank of the reference YFP among all possible variants, while the combination of T-scale with other descriptors did not improve the result (Table S5). Therefore, we used T-scale as a descriptor for our final model. The final model was trained using all measured data from the initial library including the reference YFP. This model was used to search for variants with high fluorescence performance from all possible variants in the sequence space of 20^4^.

## Acknowledgements

We thank D. A. duVerle for critical reading of our manuscript.

## Author contributions

YS conducted all the computational analysis with the support of TK. MO conducted all the molecular experiments with the support of HN and TN. KT and MU conceived of the study and directed the project. All authors participated in data interpretation and manuscript preparation.

## Data and materials availability

All data are in the paper and/or the Supplementary Materials.

## Supplementary Materials

**Fig. S1.**
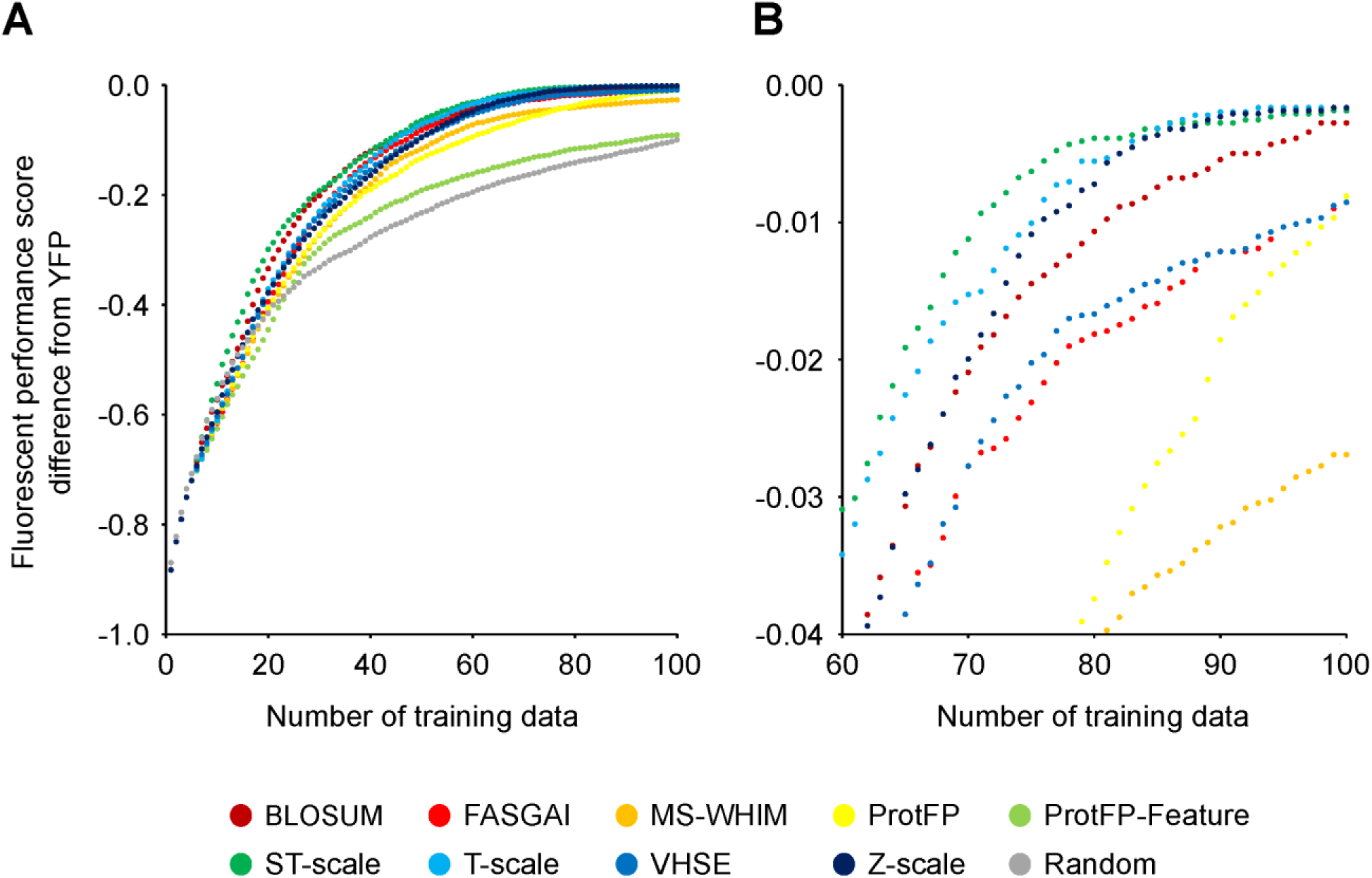
Benchmark of amino acid descriptors. For each amino acid descriptor, COMBO was trained by an Bayesian optimization procedure. The fluorescence performance score of the current best example is plotted for a given number of training data. Z-scale, T-scale, ST-scale showed better performance than the other descriptors. (**B**) is a magnified version of the late stage of training in (**A**).

**Table S1.**
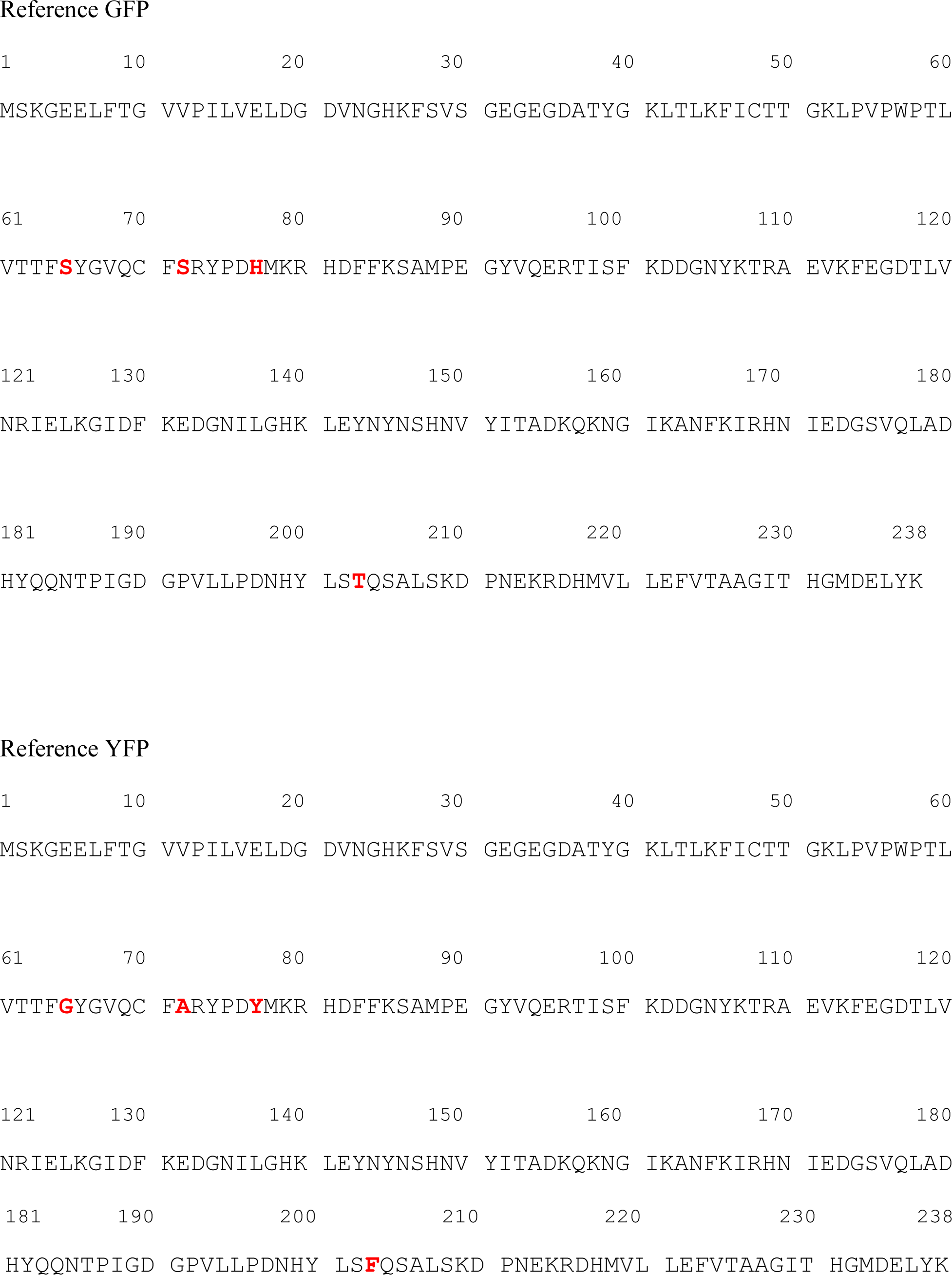
The reference GFP and YFP used in this study. The amino acid sequences are shown with the four mutations colored in red (S65G, S72A, H77Y, and T203F).

**Table S2.**
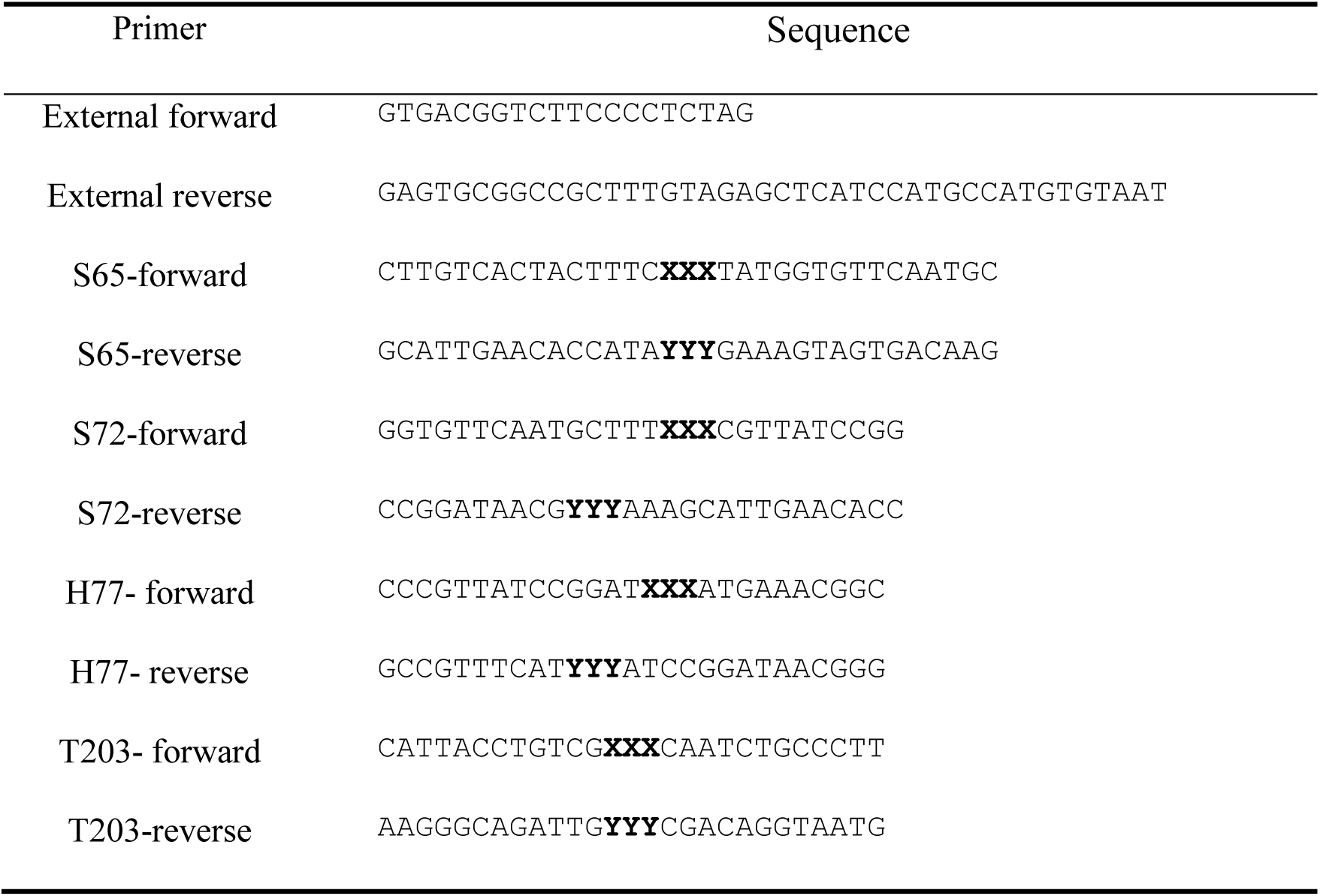
Sequences of 22-c trick primers for point saturation and site-directed random mutagenesis in GFP. XXX is NDT, VHG, or TGG. For PCR, the forward primers with NDT, VHG, and TGG was mixed at the ratio of 12:9:1, respectively. YYY is AHN, CDB, or CCA. For PCR, the forward primers with AHN, CDB, and CCA was mixed at the ratio of 12:9:1, respectively.

**Table S3.**
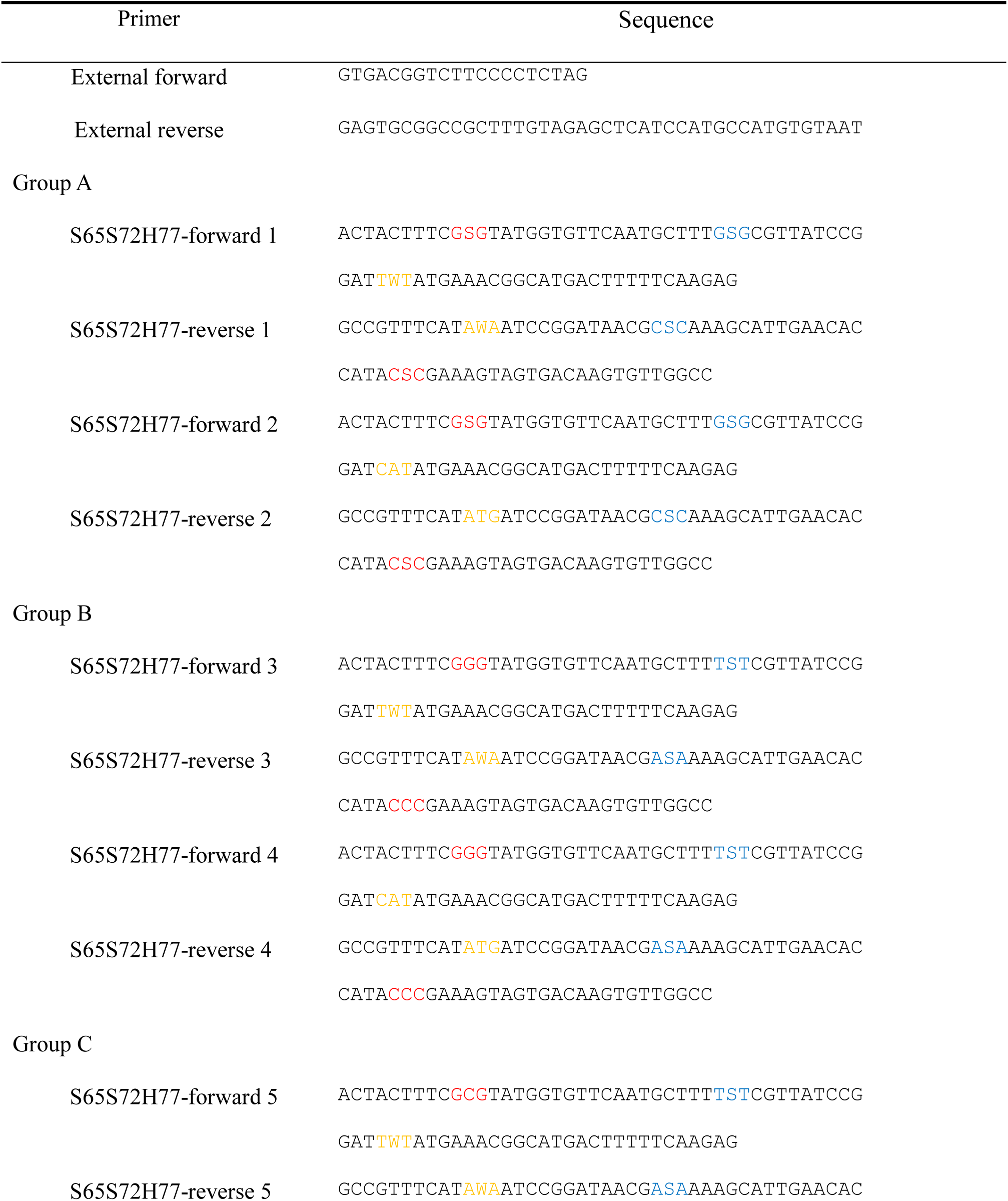
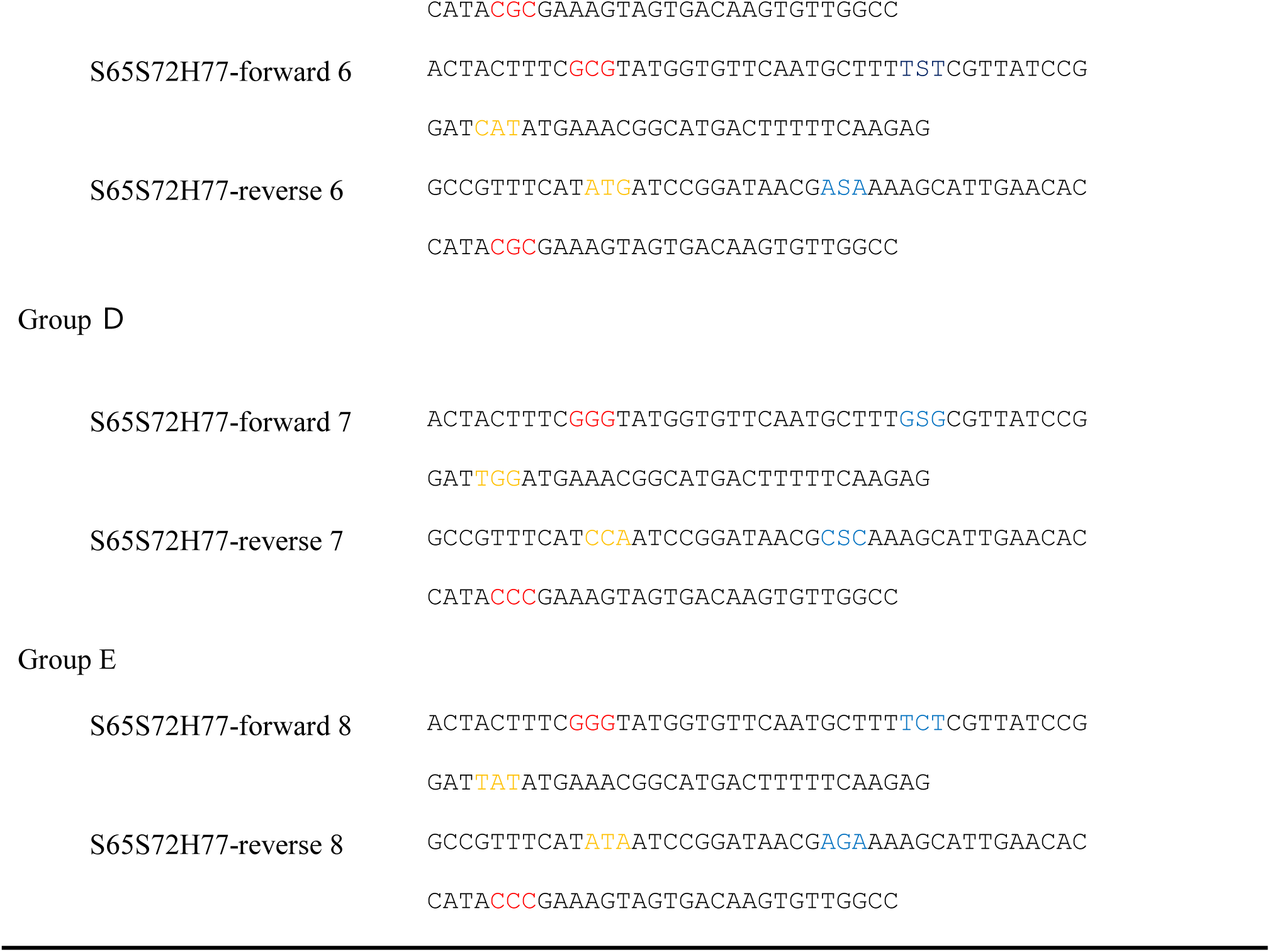
Sequences of mutagenesis primers for preparing the top 78 candidate variants proposed by machine learning. Red, blue and orange characters are the mutagenesis sites for the residues 65, 72, and 77, respectively.

**Table S4.**
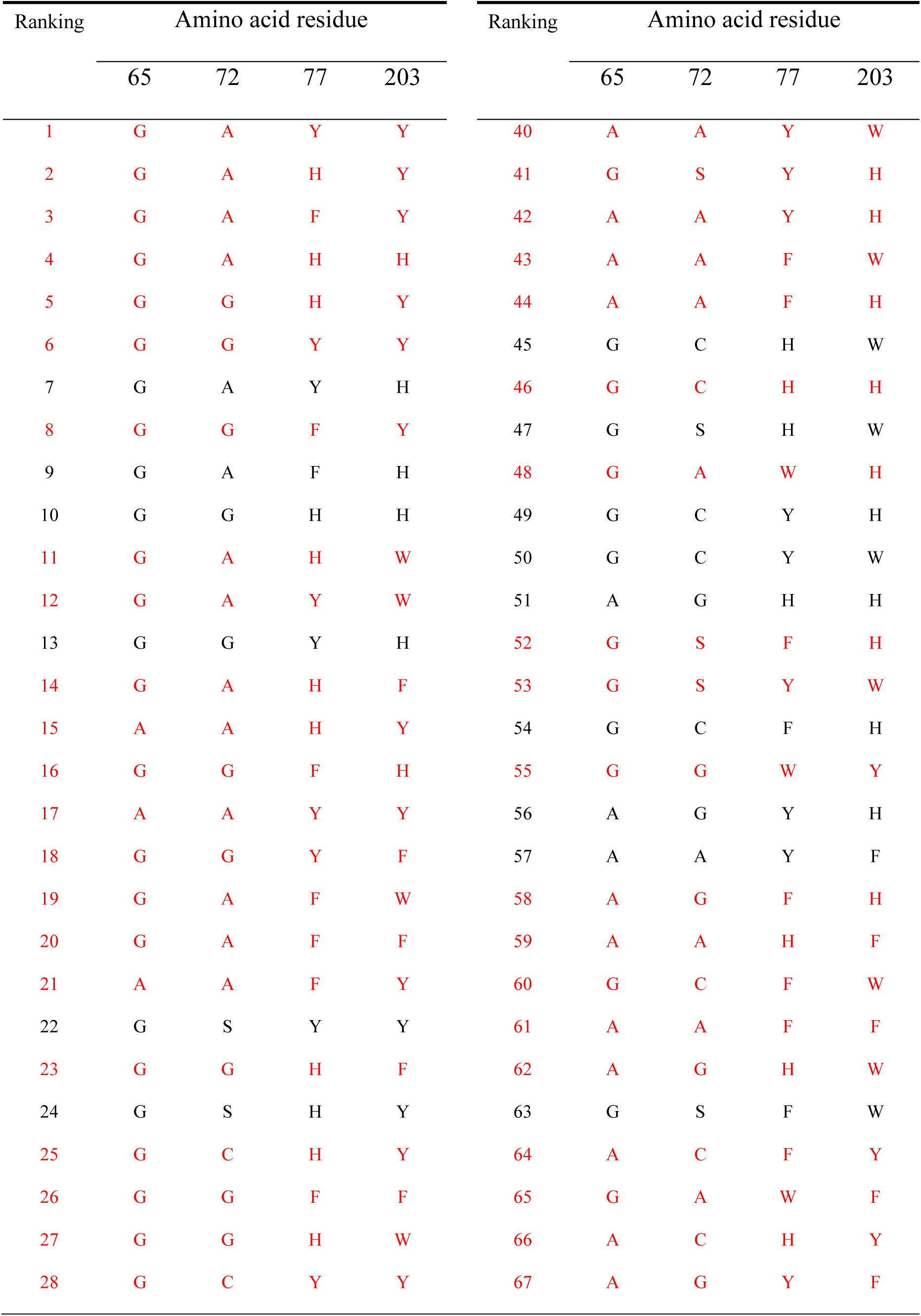
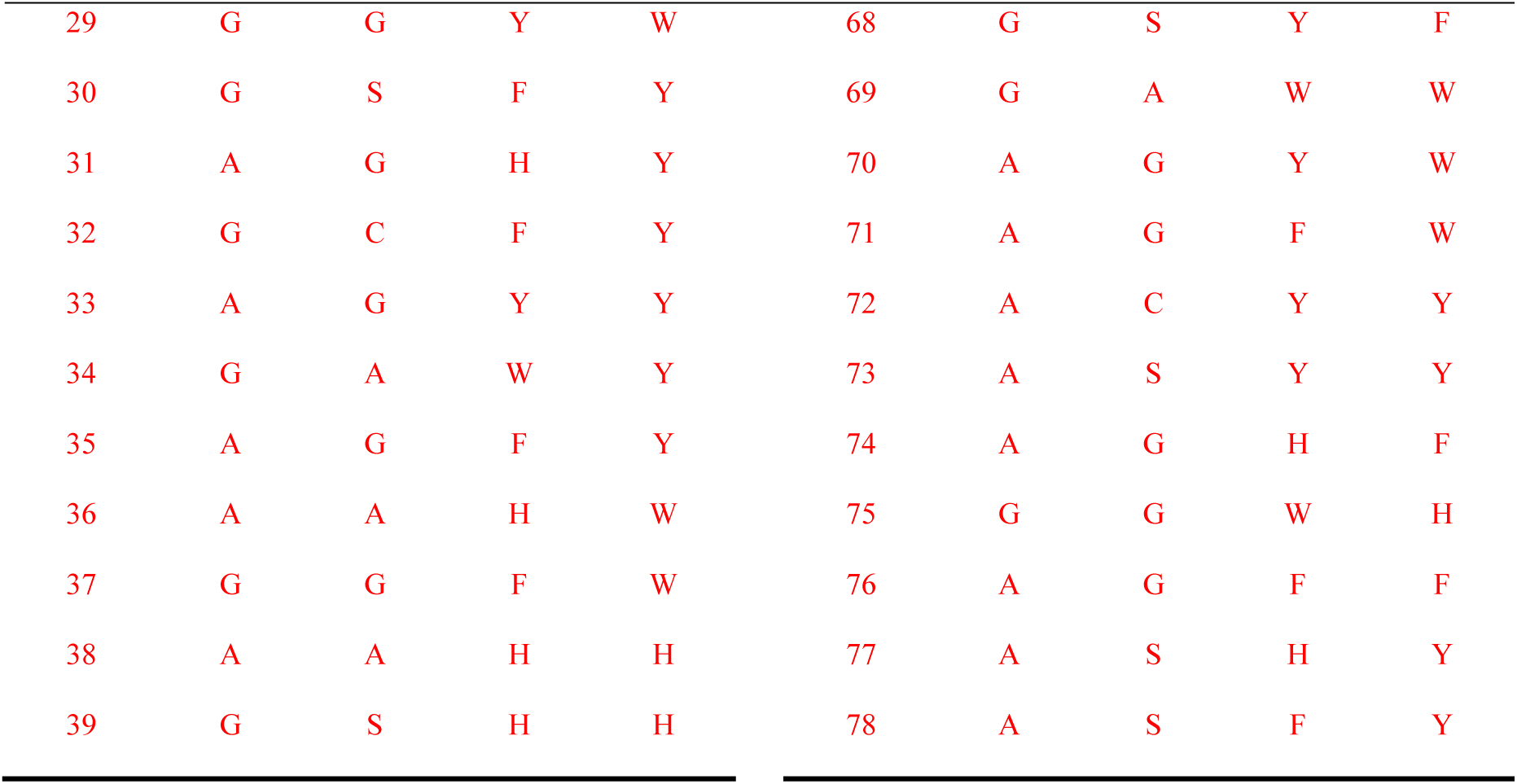
Detected GFP variants from the top 78 GFP variants. The 63 detected variants are red-colored in the table.

**Table S5.**
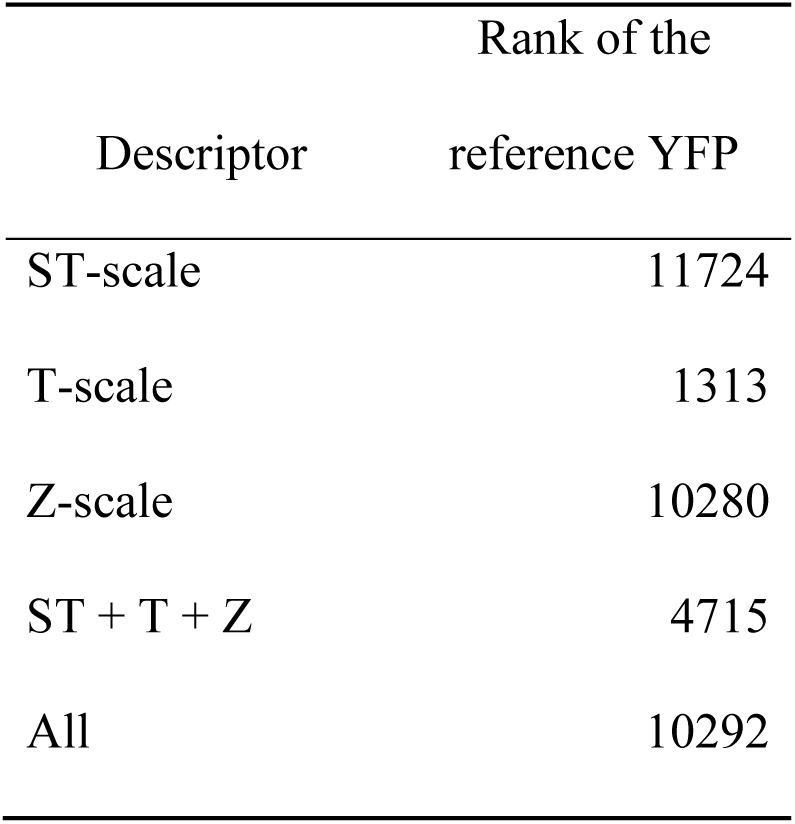
Benchmark of amino acid descriptors. ST + T + Z: feature vectors of ST-scale, T-scale, and Z-scale are concatenated into a single vector and used in machine learning. All: feature vectors of all descriptors are concatenated into a single vector and used in machine learning.

**Data file S1. Complete list of all unknown variants ranked by machine learning.**

## References

1. M. T. Reetz, L. W. Wang, M. Bocola, Directed evolution of enantioselective enzymes: iterative cycles of CASTing for probing protein-sequence space. Angew. Chem. Int. Ed. Engl. 45, 1236–1241 (2006).

2. M. T. Reetz, D. Kahakeaw, J. Sanchis, Shedding light on the efficacy of laboratory evolution based on iterative saturation mutagenesis. Mol. Biosyst. 5, 115–122 (2009).

3. M. T. Reetz, S. Prasad, J. D. Carballeira, Y. Gumulya, M. Bocola, Iterative saturation mutagenesis accelerates laboratory evolution of enzyme stereoselectivity: rigorous comparison with traditional methods. J. Am. Chem. Soc. 132, 9144–9152 (2010).

4. W. C. Lu, M. Levy, R. Kincaid, A. D. Ellington, Directed evolution of the substrate specificity of biotin ligase. Biotechnol. Bioeng. 111, 1071–1081 (2014).

5. P. J. Groot-Kormelink, S. Ferrand, N. Kelley, A. Bill, F. Freuler, P. E. Imbert, A. Marelli, N. Gerwin, L. G. Sivilotti, L. Miraglia, A. P. Orth, E. J. Oakeley, U. Schopfer, S. Siehler, High Throughput Random Mutagenesis and Single Molecule Real Time Sequencing of the Muscle Nicotinic Acetylcholine Receptor. PLoS One 11, e0163129 (2016).

6. H. A. Bunzel, X. Garrabou, M. Pott, D. Hilvert, Speeding up enzyme discovery and engineering with ultrahigh-throughput methods. Curr. Opin. Struct. Biol. 48, 149–156 (2018).

7. C. S. Karamitros, M. Konrad, Fluorescence-Activated Cell Sorting of Human l-asparaginase Mutant Libraries for Detecting Enzyme Variants with Enhanced Activity. ACS Chem. Biol. 11, 2596–2607 (2016).

8. P. Gianella, E. L. Snapp, M. Levy, An in vitro compartmentalization-based method for the selection of bond-forming enzymes from large libraries. Biotechnol. Bioeng. 113, 1647–1657 (2016).

9. A. Crameri, E. A. Whitehorn, E. Tate, W. P. Stemmer, Improved green fluorescent protein by molecular evolution using DNA shuffling. Nat. Biotechnol. 14, 315–319 (1996).

10. N. C. Shaner, G. H. Patterson, M. W. Davidson, Advances in fluorescent protein technology. J. Cell Sci. 120, 4247–4260 (2007).

11. R. Y. Tsien, The green fluorescent protein. Annu. Rev. Biochem. 67, 509–544 (1998).

12. J. S. Biteen, M. A. Thompson, N. K. Tselentis, G. R. Bowman, L. Shapiro, W. E. Moerner, Super-resolution imaging in live *Caulobacter crescentus* cells using photoswitchable EYFP. Nat. Methods 5, 947–949 (2008).

13. R. M. Dickson, A. B. Cubitt, R. Y. Tsien, W. E. Moerner, On/off blinking and switching behaviour of single molecules of green fluorescent protein. Nature 388, 355–358 (1997).

14. P. Schwille, S. Kummer, A. A. Heikal, W. E. Moerner, W. W. Webb, Fluorescence correlation spectroscopy reveals fast optical excitation-driven intramolecular dynamics of yellow fluorescent proteins. Proc. Natl. Acad. Sci. U. S. A. 97, 151–156 (2000).

15. S. Kille, C. G. Acevedo-Rocha, L. P. Parra, Z. G. Zhang, D. J. Opperman, M. T. Reetz, J. P. I. Acevedo, Reducing codon redundancy and screening effort of combinatorial protein libraries created by saturation mutagenesis. ACS Synth. Biol. 2, 83–92 (2013).

16. H. Nakazawa, R. Todokoro, Y. Ishigaki, I. Kumagai, M. Umetsu, In-one-pot-at-a-time Ligation for High-throughput Construction of a Protein Expression Vector Library. Chem. Lett. 42, 424–426 (2013).

17. F. Tian, P. Zhou, Z. Li, T-scale as a novel vector of topological descriptors for amino acids and its application in QSARs of peptides. J. Mol. Struct. 830, 106–115 (2007).

18. T. Nagai, K. Ibata, E. S. Park, M. Kubota, K. Mikoshiba, A. Miyawaki, A variant of yellow fluorescent protein with fast and efficient maturation for cell-biological applications. Nat. Biotechnol. 20, 87–90 (2002).

19. M. Kneen, J. Farinas, Y. Li, A. S. Verkman, Green fluorescent protein as a noninvasive intracellular pH indicator. Biophys. J. 74, 1591–1599 (1998).

20. B. Young, R. Wightman, R. Blanvillain, S. B. Purcel, P. Gallois, pH-sensitivity of YFP provides an intracellular indicator of programmed cell death. Plant Methods 6, 27 (2010).

21. S. Delagrave, R. E. Hawtin, C. M. Silva, M. M. Yang, D. C. Youvan, Red-shifted excitation mutants of the green fluorescent protein. Biotechnology 13, 151–154 (1995).

22. R. Heim, A. B. Cubitt, R. Y. Tsien, Improved green fluorescence. Nature 373, 663–664 (1995).

23. M. Ormö, A. B. Cubitt, K. Kallio, L. A. Gross, R. Y. Tsien, S. J. Remington, Crystal structure of the *Aequorea victoria* green fluorescent protein. Science 273, 1392–1395 (1996).

24. G. J. Palm, A. Zdanov, G. A. Gaitanaris, R. Stauber, G. N. Pavlakis, A. Wlodawer, The structural basis for spectral variations in green fluorescent protein. Nat. Struct. Biol. 4, 361–365 (1997).

25. B. P. Cormack, G. Bertram, M. Egerton, N. A. Gow, S. Falkow, A. J. Brown, Yeast-enhanced green fluorescent protein (yEGFP): a reporter of gene expression in *Candida albicans*. Microbiology 143, 303–311 (1997).

26. C. R. Stoltzfus, L. M. Barnett, M. Drobizhev, G. Wicks, A. Mikhaylov, T. E. Hughes, A. Rebane, Two-photon directed evolution of green fluorescent proteins. Sci. Rep. 5, 11968 (2015).

27. K. Sato, M. Tsuchiya, J. Saldanha, Y. Koishihara, Y. Ohsugi, T. Kishimoto, M. M. Bendig, Humanization of a mouse anti-human interleukin-6 receptor antibody comparing two methods for selecting human framework regions. Mol. Immunol. 31, 371–381 (1994).

28. T. Ueno, T. D. Rhone, Z. Hou, T. Mizoguchi, K. Tsuda, COMBO: an efficient Bayesian optimization library for materials science. Materials Discovery 4, 18–21 (2016).

29. B. Shahriari, K. Swersky, Z. Wang, R. P. Adams, N. de Freitas, Taking the human out of the loop: a review of Bayesian optimization. Proceedings of the IEEE 104, 148–175 (2016).

30. M. Sandberg, L. Eriksson, J. Jonsson, M. Sjöström, S. Wold, New chemical descriptors relevant for the design of biologically active peptides. A multivariate characterization of 87 amino acids. J. Med. Chem. 41, 2481–2491 (1998).

31. L. Yang, M. Shu, K. Ma, H. Mei, Y. Jiang, Z. Li, ST-scale as a novel amino acid descriptor and its application in QSAM of peptides and analogues. Amino Acids 38, 805–816 (2010).

32. G. Liang, Z. Li, Factor analysis scale of generalized amino acid information as the source of a new set of descriptors for elucidating the structure and activity relationships of cationic antimicrobial peptides. QSAR Comb. Sci. 26, 754–763 (2007).

33. A. Zaliani, E. Gancia, MS-WHIM scores for amino acids: a new 3D-description for peptide QSAR and QSPR studies. J. Chem. Inf. Comput. Sci. 39, 525–533 (1999).

34. G. J. van Westen, R. F. Swier, J. K. Wegner, A. P. Ijzerman, H. W. van Vlijmen, A. Bender A, Benchmarking of protein descriptor sets in proteochemometric modeling (part 1): comparative study of 13 amino acid descriptor sets. J. Cheminform. 5, 41 (2013).

35. H. Mei, Z. H. Liao, Y. Zhou, S. Z. Li, A new set of amino acid descriptors and its application in peptide QSARs. Biopolymers 80, 775–786 (2005).

36. A. G. Georgiev, Interpretable numerical descriptors of amino acid space. J. Comput. Biol. 16, 703–23 (2009).

37. M. C. Thomsen, M. Nielsen, Seq2Logo: a method for construction and visualization of amino acid binding motifs and sequence profiles including sequence weighting, pseudo counts and two-sided representation of amino acid enrichment and depletion. Nucleic Acids Res. 40, W281–287 (2012).

38. The PyMOL Molecular Graphics System, Version 2.0 Schrödinger, LLC.

